# Opsins are Phospholipid Scramblases in All Domains of Life

**DOI:** 10.1101/2025.08.17.670764

**Authors:** Zachary A. Maschmann, David E. Hardy, Indu Menon, Jenna Webb, Anant K. Menon, Katrina T. Forest

## Abstract

Opsins are highly abundant retinal proteins in the membranes of photoheterotrophic bacteria. However, some microbial genomes encode an *opsin* but lack the gene for the final enzyme in retinal synthesis. To account for this paradox, we hypothesized that bacterial opsins play a role in membrane structure and/or biogenesis independent from their potential for light-driven signaling or proton pumping. After purifying actinorhodopsin from a cell-free expression system and from *E. coli* membranes upon overexpression, we demonstrated both *in vitro* and *in silico* that actinorhodopsin from *Nanopelagicus ca.* is a phospholipid scramblase, serving in its pentameric state as a retinal-independent phospholipid diffusion channel. Phospholipid headgroups move along a transbilayer path between actinorhodopsin protomers, to equilibrate lipid content in the inner and outer leaflets. Two profound activities, membrane biosynthesis and capture of light energy, are thus facilitated by one ancient bacterial polypeptide. Light-dependent activity and light-independent phospholipid scrambling are shared functions of eukaryotic, archaeal, and bacterial rhodopsins.

**Importance:** Cells are surrounded by membranes which concentrate metabolites and protect cellular contents. Most biomembranes are phospholipid bilayers, in which the phospholipids of each leaflet orient their greasy tails inward and polar groups outward. Bilayer biogenesis depends on phospholipids synthesized on the cytofacial side of the membrane reorienting to the extracellular membrane leaflet. This reorientation requires proteins, termed scramblases, and it was shown that rhodopsins –– 7-helix photoactive membrane proteins bound to the cofactor retinal –– from organisms as widely divergent as mammals and archaea possess scramblase activity. Now we conclusively demonstrate using purified proteins in laboratory membranes as well as computational approaches, that bacterial rhodopsins are also phospholipid scramblases. This work is important because it highlights a surprising commonality among bacteria, archaea and eukaryotes and because it shows that rhodopsins – ancient proteins found in the last universal common ancestor – manifest two seemingly unrelated biochemical functions in one protein.

## Introduction

Living cells engage in an ongoing process of self-construction. Phospholipid bilayer membranes separate the cellular interior from exterior, concentrate cytoplasmic contents, and protect cellular components. Membrane production is needed for cell division, and as part of turnover and repair processes (1). Phospholipids are synthesized in the cytoplasmic leaflet of biogenic membranes, such as the cytoplasmic membrane of bacteria and the endoplasmic reticulum (2). Reorientation of phospholipids between leaflets, *lipid scrambling,* is necessary for uniform growth of the bilayer but occurs too slowly to support life if unassisted (3–8). Biogenesis of all extant membrane bilayers requires a solution to this problem (9, 10), and evolution of this function likely would have provided ancient protocells with strong fitness benefits. Indeed, compartmentalization to support early life may have relied on the modularity and renewability of lipid bilayers (11, 12). Living cells use a facilitated lipid diffusion channel, termed a scramblase, to catalyze transbilayer phospholipid equilibration (10, 13–16). Scramblases are thought to shield the charged phospholipid head group from the hydrophobic bilayer interior to expedite lipid scrambling (10, 13–16).

In recent years, rhodopsins—well known for their photon capture based activities— have been shown to moonlight as scramblases. Menon and colleagues used a fluorescence-based assay to show that purified bovine rhodopsin exhibits scramblase activity when reconstituted into large unilamellar vesicles (17), accounting for previous observations that lipids scramble rapidly across photoreceptor disc membranes (18, 19). Scramblase activity was shown to be ATP-independent and constitutive, occurring in the absence of light or the retinal chromophore. Subsequent work showed that an archaeal rhodopsin — bacteriorhodopsin (BR) from the archaeon *Halobacterium salinarum* — also had light-independent scramblase activity (20). This functional parallelism was not a foregone conclusion given that the mammalian visual sensory transducer rhodopsin and archaeal proton-pumping BR are found in anciently diverged organisms that differ greatly in membrane composition, architecture, metabolism, and environment.

Eubacterial rhodopsins have been known only since Sargasso Sea metagenomics efforts uncovered proton-pumping proteorhodopsins in marine bacteria in 2000 (21), followed by identification of opsin gene sequences in highly abundant nano-actinobacteria in fresh water (22). Like BR, these bacterial retinylidene opsins harness photon energy to pump protons across the membrane (23–25). Yet, notably, not all opsin-encoding bacteria carry genes for the complete enzymatic pathway to synthesize the retinal chromophore (23, 24). While retinal may be scavenged in some natural environments (26–28), the incomplete pathway nonetheless led us to hypothesize that actinobacterial opsins (ActR) may moonlight (29) and exhibit a second, retinal-independent function. Given that *actR* transcript levels in the natural environment are extremely high (23, 30), implying abundant protein levels, this function could be related to membrane structure and stability, echoing the fact that BR makes up 75% of the dry weight of *Halobacterium salinarum* purple membranes (31). Alternatively, given that bovine visual opsin and BR are identified scramblases (17, 20), we hypothesized that ActR and other bacterial opsins would also be phospholipid scramblases. This idea was bolstered by recent preliminary evidence of a Gram-negative bacterial proteorhodopsin (ptqPR) from *Psychroflexus torquis* acting as a scramblase (32).

We now show that ActR catalyzes interchange of phospholipid between bilayer leaflets. The dual ActR functions of proton pump (23) and scramblase emphasize the versatility of photoreceptor function and may provide a molecular connection between photoheterotrophy and membrane biogenesis.

## Results

### Large-scale synthesis of ActR in an in vitro transcription/translation system

We synthesized ActR (*Nanopelagicus ca.* genome sequence L06) in a wheat-germ extract translation system (33) with the inclusion of retinal in the translation reaction. Purification by nickel affinity and size exclusion yielded a single ActR protein band, migrating more rapidly (<25 kDa) than expected from its ∼29 kDa molecular weight, and co-purifying with a triplet of proteins identified by mass spectrometry as chaperones from the wheat germ extract (Figure 1A). The protein was highly colored (Figure 1B) with peak absorbance at 544 nm—characteristic of a bacterial rhodopsin with Leucine at the tuning position, and in agreement with our previous observations for bacterially produced ActR (23, 34) (Figure 1C). Purity and homogeneity of ActR was further demonstrated by our ability to crystallize it using a cubic lipidic phase (Figure 1B, bottom panel).

**Figure 1.**
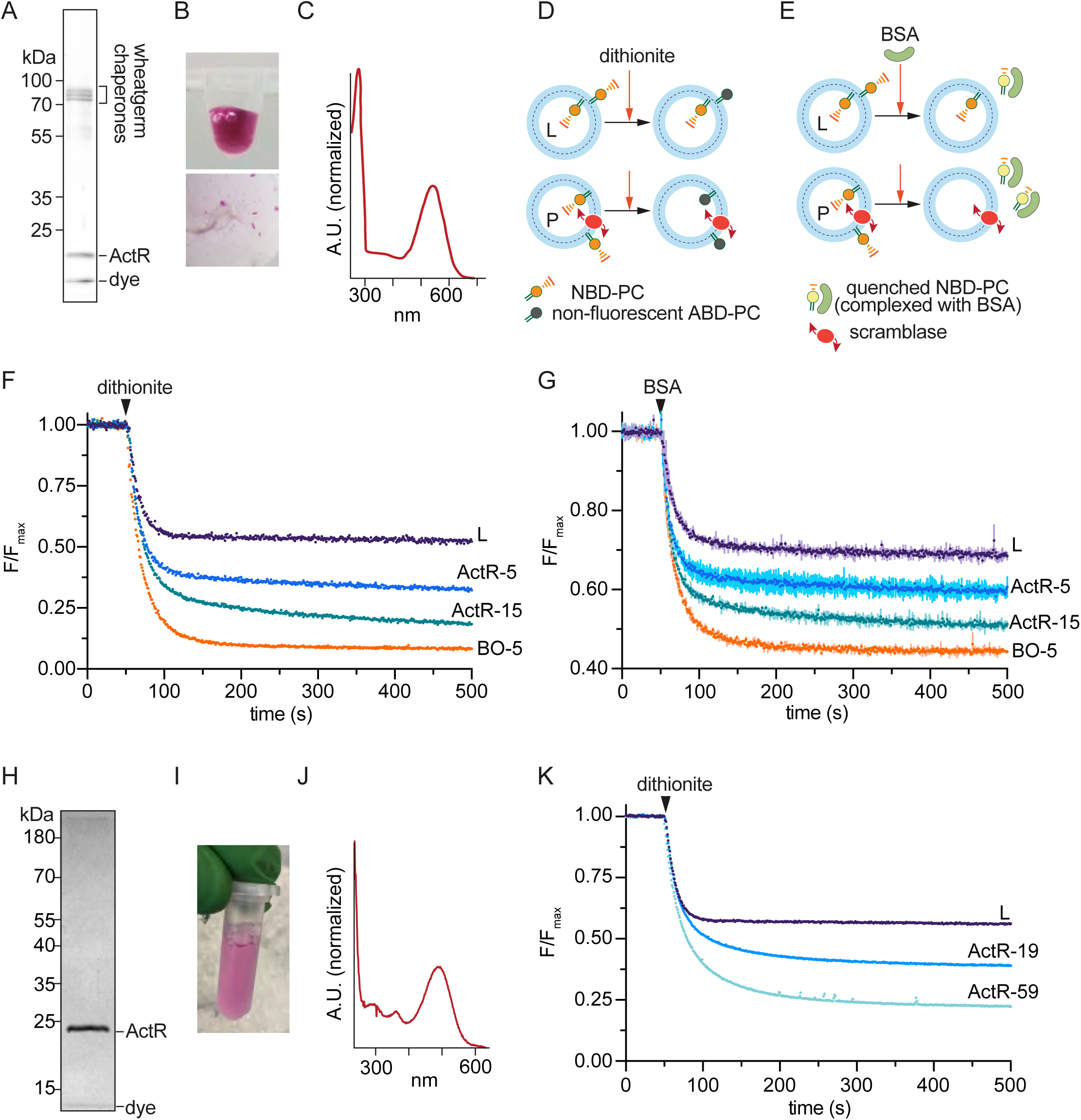
acI-B (*Nanopelagicus* ca.) ActR expressed in a cell free system or in *E. coli* is a phospholipid scramblase. (A/H) SDS-PAGE analysis of affinity-purified (A) cell free-or (H) *E. coli*-produced ActR following size-exclusion chromatography. (B/I) Image of (B) cell free-or (I) *E. coli*-produced ActR showing strong reddish color. (C/J) Absorbance spectrum of (C) cell free-or (G) *E. coli*-produced ActR. Note maximum absorbance at 543 nm. (D) Protein free liposomes (L) and scramblase-containing proteoliposomes (P) are shown schematically. The vesicles are prepared with a trace quantity of NBD-PC distributed equally between inner and outer leaflets. Upon addition of dithionite, the nitro group in NBD-PC is covalently modified to an amino group and the resulting ABD-PC is non-fluorescent. Liposomes retain fluorescence signal from NBD-PC in the inner leaflet as dithionite cannot enter the vesicles; for scramblase-containing proteoliposomes, lipids in the inner and outer leaflets exchange and fluorescence decays further as NBD-PC molecules in the inner leaflet reach the outer leaflet where they are modified by dithionite. E) As in D), except that fatty-acid free BSA is used as a topological probe. NBD-PC molecules that desorb from the outer leaflet of the outer membrane are captured by BSA. In complex with BSA, NBD-PC fluorescence is ∼60% lower than when the lipid is in the membrane. F/G/K) Normalized time course scramblase assay data for (F/G) cell free-or (K) *E. coli*-produced ActR. Dithionite or BSA was added at the indicated time point. Error bars in panel G represent standard deviation of duplicate measurements, data in panels F/G represent one of 3 independent experiments carried out under different proteoliposome reconstitution conditions, and data in panel K represent one of 4 independent experiments. Proteoliposomes were generated using ActR or bovine opsin (BO) at different protein/phospholipid ratios (units of μg/μmol) as indicated on the trace labels, e.g. ActR-5 and ActR-15 represent samples with protein/phospholipid ratios of 5 and 15, respectively.

### ActR facilitates transbilayer phospholipid transport

To determine the scramblase activity of ActR we used standard fluorescence-based assays (Figures 1D,E) in which proteins are reconstituted into large unilamellar vesicles containing a trace amount of a fluorescent phospholipid reporter, *e.g.* 4-Nitro-2,1,3-benzoxadiazole phosphatidylcholine (NBD-PC) (17, 20, 35). To assay scrambling, the vesicles are interrogated with topological probes – the dianion dithionite (Figure 1D), which reacts with the NBD fluorophore to eliminate its fluorescence, and fatty acid-free BSA (Figure 1E) which captures NBD-PC molecules that desorb from the vesicle surface, resulting in partial fluorescence quenching.

Treatment of protein-free vesicles with dithionite is expected to result in ∼50% fluorescence loss as NBD-PC molecules in the outer leaflet of the vesicles are modified whereas those in the inner leaflet are protected (Figure 1D, top row). This is what is experimentally observed (Figure 1F, trace labeled ‘L’ plateaus at 53%). However, when the vesicles contain an active scramblase, rapid exchange of NBD-PC between the inner and outer leaflets of the vesicle results in total loss of fluorescence as the entire complement of NBD-PC becomes exposed to dithionite (Figure 1D, bottom row). We reconstituted two different amounts of cell-free produced ActR via detergent-destabilization of preformed liposomes. We also reconstituted bovine opsin (BO) as a positive control. As shown in Figure 1F, dithionite treatment of ActR proteoliposomes (ActR-5, ActR-15) showed a greater loss of fluorescence than seen with empty vesicles (L), indicating that ActR has scramblase activity. Addition of hydroxylamine, which disrupts the Schiff’s base linkage to retinal, had no impact on scrambling activity (data not shown).

The reconstitutions were done at a protein:phospholipid ratio of ∼5 μg/μmol (ActR-5 and BO-5) or ∼15 μg/μmol (ActR-15). As ActR is smaller than BO (∼29 kDa vs ∼40 kDa), it would be expected to functionalize at least as many vesicles as BO when reconstituted at ∼5 μg/μmol. Yet, the data show that the extent of fluorescence reduction is not as great for ActR-5 (73%) as for BO-5 (91%) when the proteins are similarly reconstituted (Figure 1F). Even when 3x more ActR was reconstituted (ActR-15, Figure 1F), the extent of reduction was ∼84%. A possible explanation for the lower efficiency with which ActR reconstitutes is that like other opsins (20, 36–38), ActR multimerizes, and protein multimerization *en route* to reconstitution leads to a fraction of vesicles populated with trimers or monomers rather than with functional pentameric ActR scramblases as discussed below.

The fluorescence decay traces for ActR and BO-containing vesicles are well fit with a double-exponential decay function which represents a convolution of the dithionite reduction reaction rate (t_1/2_ ∼12 s) and a slower scrambling rate. For BO-5, the slow phase has t_1/2_ ∼40 s, whereas for ActR-5 and ActR-15, the t_1/2_ values are ∼400 and ∼180 s, respectively. These values suggest that ActR-mediated scrambling is somewhat slower under these experimental conditions than that observed for BO.

To complement data generated using dithionite, we next used BSA as a topological probe. For empty vesicles, ∼30% fluorescence loss is expected, as NBD-PC from the outer leaflet of the vesicles is quantitatively captured by BSA (Figure 1E, top row). This is because when bound to BSA, the quantum efficiency of NBD-PC is ∼60% lower than when the lipid is in the membrane (8, 39). For scramblase-equipped vesicles, transbilayer exchange of NBD-PC enables all NBD-PC molecules to be captured by BSA, resulting in ∼60% expected fluorescence loss. Figure 1G shows greater fluorescence loss in ActR-containing vesicles compared with empty vesicles, consistent with their scramblase activity. As with the dithionite assay, greater fluorescence reduction was seen with ActR-15 than with ActR-5, and BO-5 vesicles showed close to the maximum expected fluorescence loss.

### Scramblase activity of ActR purified from E. coli

As an orthogonal means to isolate ActR, and to avoid the co-purification of wheat germ chaperones, we overexpressed the same ActR sequence in *E. coli*, induced expression and added retinal concomitantly, and purified the protein using metal affinity and gel filtration chromatographies. SDS-PAGE analysis revealed a single band running at the expected size for an ActR monomer (Figure 1H), with a characteristic purple color (Figure 1I) and a maximum absorbance at 544 nm (Figure 1J). On reconstitution into unilamellar vesicles, the bacterially-produced protein also showed robust phospholipid scramblase activity, measured using the dithionite assay (Figure 1D, 1K). Thus, whether produced in a cell-free system or expressed in *E. coli*, ActR demonstrates scramblase activity.

### ActR oligomeric state observed by EM

Bacterial opsins have been observed in monomeric, trimeric and pentameric quaternary arrangements (37, 38, 40) as well as hexamers, although the latter may be due to native signal peptide sequences included in heterologously expressed proteins (26, 41). Extended helices and a 3-omega motif dictate the pentameric state of *Gloeobacter* rhodopsin (42). As these secondary structures are also found in ActR, we expect it to be pentameric. The multimer state of ActR may be important for both its light-dependent proton-pumping and scramblase activities. We investigated the ActR oligomeric state at the neutral pH under which the scramblase assays were carried out, using negative stain transmission electron microscopy (TEM) and analytical size-exclusion chromatography. TEM images indicate a heterogeneous mixture of ActR oligomers (Supplemental Figure 2) with pentamers identified in single particle analysis (Figure 2A). Pentamers and trimers are both present by SEC (Supplemental Figure 2), likely in equilibrium based on experimental conditions. Moreover, we observed a shift from pentamer and trimer to trimer and monomer at low pH (see further discussion in Supplemental Information). Given experimental observations and the knowledge that native functional states and most common structural assemblages of proton pumping and ion pumping bacterial rhodopsins are pentamers (26, 37, 40, 42, 43), we hypothesize that the pentameric state of ActR is the scramblase active one.

**Figure 2.**
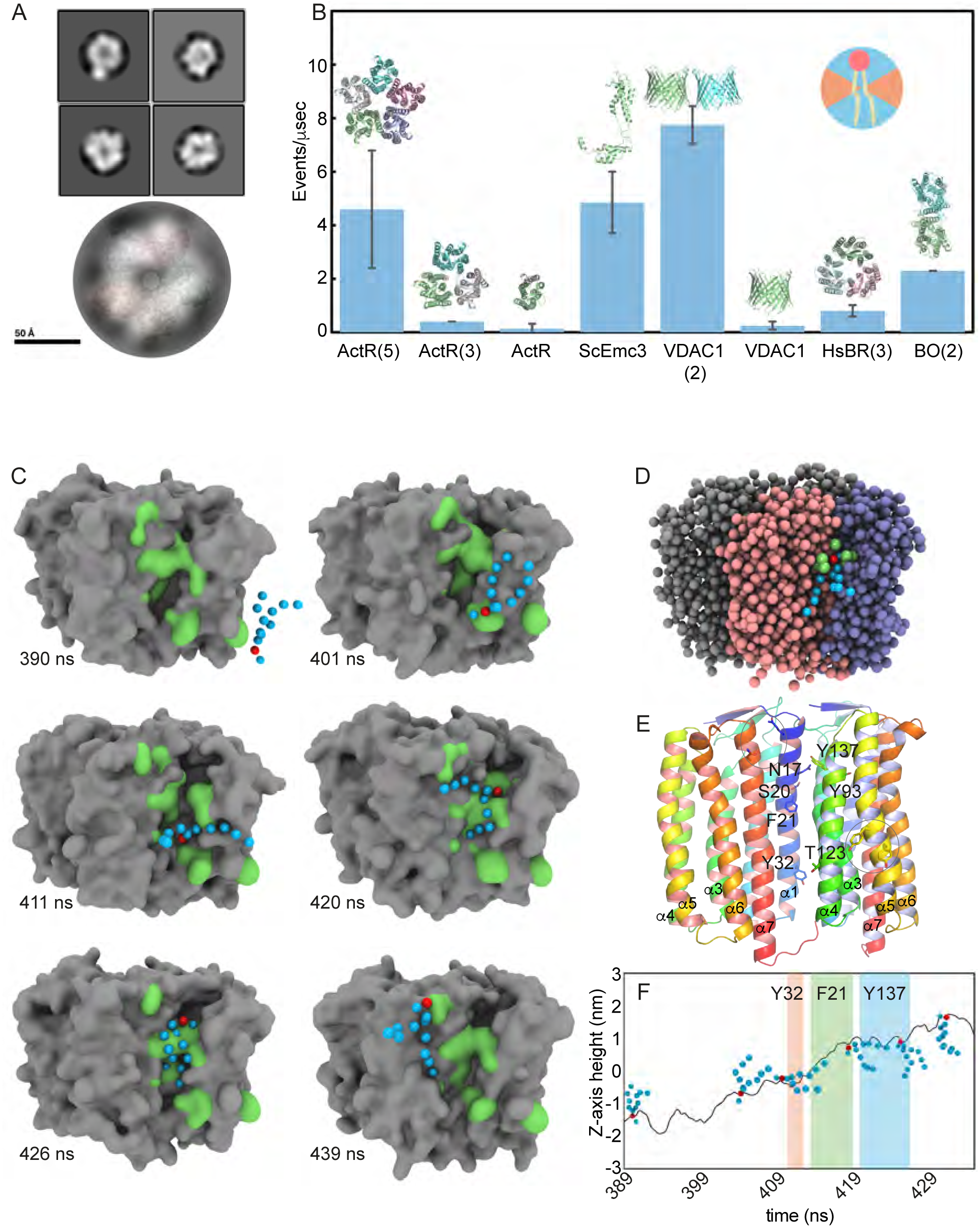
ActR is a Pentamer that Facilitates Scrambling as Assessed Computationally. A) Array of four representative negative stain electron microscopy images of ActR at pH=7.4, and superposition of representative ActR TEM image over predicted ActR pentameric structure. B) CGMD scramblase events/μsecond accompanied by structural images. The upper right inset shows the angles (in orange) in which rotations were not counted as scrambling events. C) CGMD scramblase “flip book” images of one DOPC moving through cleft and re-orienting lipid tails. Those beads which were within 5 Å of the phosphate bead in any frame are shown in green, PO4 bead is red, and lipid tail beads are blue. D) One snapshot from the trajectory represented in part C is shown with beads from each opsin monomer shaded differently to highlight the inter-protomer location of the groove. Green protein beads are all those within 5 Å of the red phosphate bead (in this frame representing atoms from N17, S20, F21 and Y93), while blue beads depict the lipid tail. E) Ribbon diagram of two adjacent protomers that form the scrambling groove, with side chains in the groove that interact with lipids during scrambling labeled, and occasionally observed interactions circled but not labeled. Each monomer is colored in a reverse rainbow from blue –> red as the chain progresses from N –> C terminus. Additionally, the left chain has a salmon interior surface while the right chain has a light blue interior surface, with visible α helices labeled on both chains. F) Z-axis height above or below the center of the membrane for the PO4 bead as a function of time during passage through the cleft of the example individual DOPC shown in panel C. Dwell times for Y32, F21, and Y137 depicted as colored bars.

### Coarse-grained molecular dynamics support ActR pentamer scramblase activity

To support our laboratory-based observations that ActR is a lipid scramblase, we employed coarse-grained molecular dynamics (CGMD) simulations over an extended time window to assess scrambling activity of ActR in three oligomeric states. We used the computational protocol of Li *et al.* (44) to determine the scrambling rate by counting the number of phospholipid inversions in the membrane per unit time. Three replicates of an ActR pentamer model in a dioleoylphosphatidylcholine (DOPC) membrane bilayer showed ∼4 scrambling events/µs (Figure 2B), consistent with rates calculated for archaeal BR (20).

The lipid inversion rate of two positive controls (scEMC3 with ∼5 events/µs and VDAC1 dimer with ∼8 events/µs) were consistent with results reported by Li *et al.* (44). In contrast, the ActR monomer, ActR trimer, and VDAC1 monomer displayed negligible scrambling activity (Figure 2B).

Lipid translocation by the ActR pentamer shows no directional bias and, although durations vary, on average the complete inversion takes on the order of 100 ns. Our model suggests that transbilayer phospholipid diffusion occurs at protomer interfaces in ActR and involves polar and aromatic residues that form a hydrophilic groove, consistent with the previously described credit card model (15). Side chains spanning the hydrophilic groove derive from α-helix 1 (α1) of one protomer and α3 and α4 of the adjacent protomer (Figure 2C,D,E).

The scrambling trajectory can be described in steps. First, lipids are recruited by polar residues near the bilayer surface. For example, Tyr161 and Tyr164 on α5 serve this role in the event depicted in Figure 2C. The phospholipid does not linger in these positions for more than a frame or two during the CGMD simulation, and from 20 scrambling events chosen for a deeper analysis (of total 138 observed), these residues come into play in only 1 or 2 of the events, respectively. Thus, the recruitment step is transient and many different side chains may be able to act at this step.

After this initial recruitment of the phospholipid, the heart of the scrambling mechanism takes place within the inter-subunit groove. The phosphate bead enters a lower pocket and contacts Tyr32 (75% of events) and Thr123 (95% of events) (Figure 2C,E). During this step, the lipid tails may sample an equatorial orientation relative to the bulk lipid of the membrane (Figure 2C, 411 ns & 420 ns panels). Subsequently, Phe21 (100% of 20 events) passes the DOPC molecule from the lower to an upper pocket. Passage into and through this second water-containing cavity formed by Asn17, Ser20, Tyr93 and Tyr137 (contacted in 90%, 95%, 40% and 85% of events, respectively) is slow, with variable contact times on the order of 10 ns for these residues (Figure 2C-F). The buried water molecules are displaced from this major pocket when the PO4 bead enters. All of these transitions can include backsliding. The upper cavity is formed by side chains from three helices at the oligomer interface; Asn17 on α1 of one monomer interacts with Tyr93 on α3 and Tyr137 of α4 from the adjacent monomer (Figure 2C,E).

Finally, an additional step involves transient contact with surface residues (for example, Gln11 and Asn220), after which the lipid diffuses into the bulk membrane.

### Retinal does not aXect scrambling in silico but polar residue substitutions along cleft do

ActR scramblase activity is not changed in laboratory experiments with the addition of hydroxylamine, nor was illumination part of our protocol. Retinal was not present in our CGMD simulations. Thus, scramblase activity of ActR does not depend on chromophore or light. Nonetheless, to assess whether retinal has an effect on scrambling rates *in silico*, we built a simplified bead model of retinal and included it in additional computational tests of scramblase activity (Supplemental Figure 2). The scrambling rate was not markedly different in these MD simulations from those without retinal.

In order to computationally evaluate the importance of the α1 and α4 polar amino acids as core elements of the scrambling groove, we created four ActR variants with two substitutions each and tested their scrambling activity as described above for the native sequence. Consistent with the identification of polar residues in the cleft as major contact points for the PO4 bead, both our double substitutions N17L/Y137F and N17L/T123L showed a reduced scrambling of ∼1 event/µs (Supplemental Figure 2).

## Discussion

Moonlighting is common among identified scramblases — multiple scrambling GPCRs (36, 45) and TMEM16 (46) in eukaryotes, numerous bacterial, ER, and mitochondrial membrane insertases (44, 47), and Gram positive (this work), Gram negative (32) and archaeal rhodopsins (20) perform other functions including ion transport, protein translocation, and signal transduction. Members of the TMEM16 family of proteins, for example, function either as chloride channels or as nonspecific ion channels/lipid scramblases. The chloride channel members of this family may have derived from a primordial scramblase and evolved chloride transport at the expense of scramblase activity (48). It appears likely that yet-to-be discovered scramblases moonlight with other functions. The ubiquitous nature of scramblase activity in diverse membrane proteins also helps explain why it has been difficult to pinpoint a unique and essential scramblase in some organisms (5).

Rhodopsins were ancient proteins found in early-Earth life forms (49, 50), suggesting they could have functioned as scramblases in the earliest cells. In the modern world, microbial rhodopsins are found across Bacteria, Archaea, and even some giant viruses (51, 52), with phylogenetic evidence suggesting that this wide distribution is due, at least in part, to frequent horizontal gene transfer (53). We are not aware of any conserved sequence motifs linked to the scramblase functionality of rhodopsins across the domains of life. This lack of sequence requirement suggests that scramblase functionality relies upon relatively hydrophilic surface characteristics without selecting for specific residues. Moreover, we do not expect opsin scramblase activity to be specific with respect to lipid head groups, as known scramblases are promiscuous for their substrates but do not have unlimited specificity for complex head groups (17, 54, 55). While scramblase activity is essential, the molecular surface underpinning this function appears to be under low selective pressure, possibly owing to the high rates of lipid scrambling exhibited by even rudimentary scramblases (56, 57) that may render scramblases easily evolvable.

Our simulations show that the ActR pentamer exhibits robust scrambling activity, consistent with our experiments showing that more ActR is needed to functionalize vesicles in comparison to bovine opsin. As monomeric and trimeric forms of ActR showed nominal scrambling, a fully functional, scrambling-competent hydrophilic groove is only present in the pentameric form. The dependence of activity on oligomerization aligns with observations from other scramblases such as VDAC1, where dimerization is necessary to form a translocation-competent groove (39) and BR, where trimerization generates interfaces that can scramble lipids (20).

The structural basis for actinorhodopsin-mediated scrambling centers on a hydrophilic groove formed by helices 1 and 3+4 of neighboring protomers. This pathway is lined with aromatic and polar residues that interact with the charged phospholipid headgroup to facilitate translocation. The orientation of the phospholipid during the scrambling event supports the credit-card insertion model, where the charged phosphate head group slides between helices, forming transient interactions with polar groups, and reorients in plane with the bulk lipid as it rejoins the bilayer (Figure 2C,D,F). Coarse-grained resolution limits the precision with which we can observe side chain orientation and hydrogen bonding. However, repeated observations of lipid contacts indicate a core set of residues—including Asn17, Phe21, Tyr32, and Tyr137—that play a role in the translocation mechanism.

We have shown that ActR from *Nanopelagicus ca.* of the acI lineage of actinobacteria is a phospholipid scramblase, making it the third known microbial opsin with scramblase activity alongside Gram-negative bacterial PtqPR and archaeal BR. Our demonstration of the robust scrambling activity of Gram-positive bacterial actinorhodopsins supports the notion that phospholipid scrambling is a fundamental function of rhodopsins across the domains of life. This insight provides a broad foundation for further study of the scramblase/opsin duality. Particularly intriguing open questions are 1) Is there a pH dependence of scrambling activity? The oligomeric state of *Gloeobacter* rhodopsin has been shown to vary with pH (38) and our size exclusion chromatography mirrors these results (Supplemental Figure 2). Moreover, our computational results show that trimers don’t scramble (Figure 2B), thus providing a consistent model that the ActR low-pH oligomer is not an active scramblase. Decreased scramblase activity at low pH would prevent dissipation of the proton gradient; low phospholipid pKa values enable a scenario where outer leaflet phospholipids are protonated and subsequently deprotonated when scrambled to the inner leaflet (58). Though *Nanopelagicus ca.* primarily inhabits the fresh water epilimnion, a neutral pH habitat, freshwater lakes and bogs can contain low pH zones that may provide environmental pressure for a pH-dependent adaptation of scramblase activity by ActR (59). Perhaps this is a common mechanism for regulating bacterial rhodopsin scramblase activity with consequences for energetics as well as membrane biogenesis. 2) Does the secondary antenna carotenoid associated with rhodopsin in many bacterial and archaeal opsins (23, 60–62) have an impact on scrambling? Further elucidation of the structural roles of secondary carotenoids as well as scrambling mechanisms is necessary to understand the interplay, if any, between antenna association and scramblase functionality. Although carotenoid association may add to thermal stability (61), our experimental and computational scrambling assays were all done in the absence of the actinorhodopsin carotenoid (23) and thus we can confidently say the antenna is not required for scramblase activity. 3) Is the earliest role of rhodopsin scrambling or photoheterotrophy, or did these activities co-evolve? Using our CGMD pipeline, we evaluated the scramblase activity of a plausible ancient opsin sequence recently proposed by ancestral sequence reconstruction (49). We did not observe scramblase activity (data not shown). While this model has very highly conserved residues associated with photoactivity, our current study concludes that scramblase activity relies more subtly on appropriate alignment of amino acids with relevant properties along a protomer interface than on invariant side chain identities. The study of the evolution of scramblase activity in early opsins is an exciting area that warrants additional analysis.

## Materials and Methods

### Materials

16:0-18:1 PC (POPC) (1-palmitoyl-2-oleoyl-glycero-3-phosphocholine), 16:0-18:1 PG (POPG) (1-palmitoyl-2-oleoyl-sn-glycero-3-phospho-(1’-rac-glycerol) (sodium salt)), 16:0-06:0 NBD PC (C6-NBD-PC) (1-palmitoyl-2-{6-[(7-nitro-2-1,3-benzoxadiazol-4-yl)amino]hexanoyl}-sn-glycero-3-phosphocholine) and 14:0-06:0 NBD PC ((1-myristoyl-2-{6-[(7-nitro-2-1,3-benzoxadiazol-4-yl)amino]hexanoyl}-sn-glycero-3-phosphocholine) were purchased as chloroform solutions from Avanti® Polar Lipids. n-Dodecyl-B-D-maltoside (DDM) was purchased from VWR International. Bio-Beads SM2 Adsorbents were purchased from Bio-Rad. Amberlite XAD-2 and sodium dithionite were purchased from Sigma Aldrich, and fatty acid-free BSA was from EMD Millipore. Gibson Assembly® Master Mix was purchased from New England BioLabs. PhusionTM High-Fidelity DNA Polymerase Transcription reaction components, including S6 RNA polymerase, NTPs, RNase inhibitor, and buffer were purchased from Promega Corporation. Translation reaction components, including WEPRO® 8240H, Creatine Kinase, and SUB-Amix, were purchased from CellFree Sciences.

### Plasmid Construction

DNA codon-optimized for expression in *E. coli* and encoding the amino acid sequence of ActR (ActR_L06_ from *Nanopelagicus* ca. single-cell amplified genome L06 (acI-B)) (63, 23) was cloned into a pEU-C-His flexiVector, chosen for its suitability for *in vitro* protein production, under the transcriptional control of the SP6 promoter (33). Insertion of the coding sequence for ActR extended with a C-terminal 6x-His tag into the pEU flexiVector was performed by amplifying plasmid fragments using PCR followed by Gibson Assembly (64, 65). The same DNA sequence was also inserted behind the T7 promoter of a pET-28a vector using Gibson Assembly for expression in *E. coli*. Amplification of DNA fragments was performed using Phusion^TM^ High-Fidelity DNA Polymerase. Gibson Assembly was performed using Gibson Assembly^®^ Master Mix. The molecular weight of the 6x-His-tagged ActR monomer is 29.7 kDa, with a predicted pI of 8.64.

### Large Scale Cell Free ActR Production

Large scale transcription and translation reactions were carried out essentially as described in Aly *et al.* (33, 66) using low magnesium buffer (Cell Free Sciences) for transcription. Briefly, ActR-His_6_ protein was synthesized in a 12-ml 8240H wheat germ extract cell-free translation system (Cell Free Sciences) at 60 OD, using 0.1% MNG-10 (maltose neopentyl glycol-10) with 0.02% cholesteryl hemisuccinate (CHS) buffer, and 0.1 mM all-trans retinal was added during the translation. ActR was purified by Ni-affinity chromatography followed by size exclusion. Protein was concentrated to approximately 15 mg/ml in 10 mM Hepes pH 7.5, 100 mM NaCl, 0.3 mM TCEP, 0.05% DDM, 0.01% CHS and stored at –80 C. The contaminating higher molecular weight bands were identified by mass spectrometry to contain uncharacterized wheat protein with Uniprot ID BAK08004.1, NADH Dehydrogenase, and HSP 90.

### ActR Expression and Purification in E. coli

BL21 (DE3) *E. coli* cells were used for recombinant expression of ActR. 10 mL of an overnight culture was used to inoculate 1 L of LB supplemented with 50 μg/mL kanamycin, which was then incubated with shaking (37°C, 250 rpm) until the OD_600nm_ reached 0.3, at which point the temperature was reduced to 22°C. At OD_600nm_ of 0.5-0.6, ActR expression was induced by adding powdered IPTG to a final concentration of 0.5 mM. All-trans retinal was added concomitantly to a final concentration of 9 μM from a 50 mg/mL stock solution in ethanol. After 16-20 hrs of growth in the dark, the culture was centrifuged at 4,000 g for 12 minutes at 4°C. After a freeze-thaw cycle, cells were resuspended in lysis buffer (50 mM Tris, 300 mM NaCl, 5 mM MgCl_2_, 130 μM CaCl_2_, 1 mg/mL lysozyme, 1 mM PMSF, pH=7.40), and sonicated. This and all subsequent steps were carried out at 4°C. The sample was clarified at 10,000g for 20 min, after which the supernatant was centrifuged at 40,000g for 45 min to pellet membranes. The membrane fraction was resuspended in 20 mL of solubilization buffer (50 mM Tris, 300 mM NaCl, 5 mM MgCl_2_, 130 μM CaCl_2_, 10 mg/mL DDM, pH=7.40) using a Potter homogenizer, and incubated with rocking in the dark for 16-20 hrs. Aggregates were removed by centrifugation at 40,000g for 20 min. The supernatant (∼20 mL), now containing solubilized membrane proteins, was collected and flowed continuously over 5 mL of Ni-NTA resin at 1 mL/min for 16-20 hrs using a peristaltic pump. The Ni-NTA resin was washed with >20 bed volumes of wash buffer (50 mM Tris, 300 mM NaCl, 50 mM Imidazole, 0.5 mg/mL DDM, pH=7.40) and ActR was eluted with wash buffer containing successively higher imidazole concentrations, from 100 mM to 500 mM. ActR-containing fractions were pooled, and concentrated along with DDM in a 10 kDa – cutoff spin concentrator (Millipore). Samples were dialyzed against 50 mM HEPES, 150 mM NaCl, pH=7.4 and injected onto a Superdex 200 Increase 10/300 GL size exclusion column pre-equilibrated with buffer identical to the dialysis reservoirs supplemented with 0.5 mg/ml DDM using an ӒKTA pure™ chromatography system. ActR-containing fractions were pooled, concentrated to approx. 0.5 mg/ml and immediately used for liposome reconstitution. The overall yield was ∼0.5 mg/L of bacterial culture.

### ActR Negative Stain EM

ActR expressed in *E. coli* and purified through the metal affinity chromatography step was kept at 4°C and used to prepare EM grids on the same day. CF200-CU grids were glow discharged for 30 s at 15 mA on a Pelco easiGlow discharge system. 4 µL of sample (diluted in elution buffer to 240, 80, 32, or 8 µg/mL) was applied to each grid for 1 minute before blotting. Grids were washed 2x in 20 µL buffer, 2x in 20 µL 2% Uranyl acetate (UA), followed by one minute incubation in 2% UA. All images were collected on a Talos L120c 120 keV electron microscope using a Ceta CMOS detector at the University of Wisconsin–Madison Cryo-EM Research Center.

### Reconstitution of ActR into liposomes

Liposomes were prepared using a 9:1 (mol/mol) POPC/POPG mixture, spiked with 0.5% 16:0-06:0 NBD PC or 14:0-06:0 NBD PC. The lipid mixture was prepared from chloroform stock solutions, dried under nitrogen or in a rotary evaporator, and then placed under vacuum overnight at ambient temperature before being resuspended in buffer (50 mM HEPES, 150 mM NaCl, pH 7.4) at a concentration of ∼4 mg/mL. The resuspended lipids were processed further and taken for ActR reconstitution using either of two methods as follows. Method 1 (used for *E. coli*-produced ActR): The sample was bath sonicated for 5 min, incubated at ambient temperature for 1 h, sonicated again and aliquoted for individual reconstitution experiments. Each aliquot was incubated with DDM (0.5% final) at ambient temperature for 1 h, rotating end-over-end. ActR was then added, and incubation continued for 1 h. Reconstitution was achieved by adding washed Amberlite XAD-2 (150 mg) followed by incubation at ambient temperature for 1 h with end-over-end rotation. The samples were withdrawn and added to a fresh aliquot of Amberlite, followed by incubation overnight at 4°C with end-over-end rotation. Following this, the samples were withdrawn and transferred to tubes containing fresh Amberlite for a final incubation at ambient temperature for 1 h. The resulting proteoliposomes were stored at 4°C for up to two days, diluted 1/10 in excess buffer and passed through a 0.2 μm filter on a Hamilton syringe assembly to produce a narrow size distribution of ActR-containing liposomes (67, 68). Method 2 (used for cell-free synthesized ActR): The sample was extruded successively through 400 nm and then 200 nm filters as described (39) to generate unilamellar vesicles with an average diameter of 170 nm. TX-100 (0.2% final) was added to destabilize the vesicles, before adding ActR. The mixture was treated with one shot of washed BioBeads to remove detergent, initially at room temperature and then overnight at 4°C. The resulting vesicles were assayed directly (no further extrusion).

### Scramblase Assays

Scramblase activity assays were performed as previously described (17, 20, 69, 70), using vesicles diluted to 2.5 mL in buffer (50 mM HEPES, 150 mM NaCl, pH7.4). NBD fluorescence (λ_ex_=470 nm, λ_em_=530 nm) was monitored over time in spectrofluorimeters (QuantaMaster Model C-60/2000 or Horiba) equipped with a temperature-controlled cuvette holder, with a magnetic stirring apparatus. After obtaining a stable fluorescence signal, dithionite (from a 1 M stock) or fatty acid-free BSA (from a 75 mg/ml stock) was added. Addition of dithionite (final 10-20 mM) covalently modifies the fluorescent NBD moiety and eliminates its fluorescence signal (71), leading to fluorescence drops of 50-100% for protein-free liposomes and scramblase-active proteoliposomes, respectively. BSA (final 15 mg/ml) captures and quenches NBD-PC molecules that desorb from the outer leaflet, leading to fluorescence drops of ∼30-60% for protein-free liposomes and scramblase-active proteoliposomes, respectively (8, 72).

Fluorescence traces were normalized to stable fluorescence emission values prior to dithionite or BSA addition, and analyzed as described (73) using mono-exponential fits for protein-free liposomes and a double-exponential fit for ActR-containing samples. Analyses were performed within MATLAB or GraphPad Prism, using their curve fitting functions.

### CGMD

An ActR monomer structural model was generated using Chai-1 for *de novo* structure prediction (74). The crystal structure of VDAC1 (RCSB: <u>6G6U</u>) and bovine opsin (RCSB: 4A4M) were obtained from the RCSB Protein Data Bank (75). For scEMC3, missing regions in the crystal structure (<u>6WB9</u>) were modeled using AlphaFold3 (76, 77). Two BR dimers were tested and compared; one generated in AlphaFold3 and aligned to a crystal structure (4MD2) and the other a vacuum-minimized crystal structure (1AT9). We found similar scramblase activity in both models, and herein we present the AI-predicted model. CGMD simulations were performed on a high-throughput computing cluster using the HTCondor software suite and GROMACS 2022.2 (78, 79). Visual Molecular Dynamics (VMD) and Tachyon were used for trajectory visualization and image rendering (80, 81).

Protein models were initially subjected to energy minimization in a vacuum until the maximum force on any atom was below 100 kJ·mol⁻¹·nm⁻¹. Both the minimization and production runs were conducted using a GROMACS 2022.2 container image hosted on Docker Hub. Following minimization, the ActR trimer and pentamer were built by aligning the monomer to a trimeric *Halobacter salinarum* BR model (<u>1AT9</u>) (82) and to a pentameric sodium-pumping rhodopsin (KR2) from *Dokdonia eikasta* (<u>6YC3</u>) which was crystallographically determined at 2.1 Å resolution (37). VDAC1 monomers were aligned to the anti-parallel dimer RCSB structure and subsequently manually adjusted to the highly scramblase-active parallel conformation described by Jahn et al. (39).

Proteins were embedded in a dummy membrane using the Orientation of Proteins in Membranes (OPM) web server (83). The resulting protein–membrane systems were built and solvated using CHARMM-GUI, then converted to coarse-grained representations using the Martini 3 force field with Elastic Network Dynamics (ElNeDyn) (84–88). Each membrane leaflet contained 400–500 molecules of 1,2-dioleoyl-sn-glycero-3-phosphocholine (DOPC), for a total of 800–1000 lipid molecules per bilayer. Electrostatic charges in the protein were counterbalanced by adding Na⁺ and Cl⁻ ions to solution to bring the net charge to zero. Each system was solvated with approximately 2 × 10^4^ water beads.

To profile lipid dynamics in the presence of putative scramblases, simulations consisted of energy minimization, equilibration, and a production run with randomized initial velocities. Production simulations were run for 5 × 10^8^ steps with a time step of 20 fs, yielding a total trajectory length of 10 µs. Temperature was maintained at 310 K using a velocity-rescaling (v-rescale) thermostat within an NPT ensemble. Pressure was set to 1 bar using the Parrinello–Rahman barostat. Calculations were executed on a local GROMACS container and using HTCondor at the UW-Madison Center for High Throughput Computing (89).

Systems were interrogated for scramblase activity by measuring phosphate headgroup (PO_4_) angles of every DOPC every 1 ns following the method described by Li *et al*. (44). A vector is defined whose direction bisects the angle between the two vectors from head group to lipid tail for each DOPC. A buffer region is defined from 55° to 125° and an event is measured if the vector goes from under 55° to over 125° (or *vice versa*). Denoising was accomplished by averaging angles over 50 ns windows and sampling this smoothed data every 25 ns. The number of scrambling events per microsecond were averaged across all replicates. Simulations for VDAC1, scEMC3, BO, and ActR trimer were run twice and all other models were run three times.

## Acknowledgements

We thank Prof. Brian Fox and Dr. Emily Beebe of the UW-Madison Transmembrane Protein Center for *in vitro* production and purification of ActR, Dr. Takefumi Morizumi and Prof. Oliver Ernst (University of Toronto) for a sample of bovine opsin, and Prof. Katherine Henzler-Wildman, Dr. Vilius Kurauskas and Prof. Katherine McMahon for valuable expertise. We thank the Landick and Kaçar labs at the University of Wisconsin–Madison for use of laboratory equipment. We thank Prof. Elizabeth R. Wright, Dr. Bryan Sibert, and Alex Cotton at the University of Wisconsin–Madison Cryo-EM Research Center for negative stain cryo-EM services.

## Funding and Additional Information

Research was supported by the USDA (WIS03052 to KTF), Einstein Foundation Visiting Fellow Program (to KTF), the Human Frontier Science Program (Ref.-No: RGP009/2023 to AKM) and the National Science Foundation (Grant 2425586 to AKM and Graduate Research Fellowship Grant No. 2137424 to DEH). The Transmembrane Protein Center was funded by the NIH (1U54 GM094584 to Brian Fox). Some of the fluorescence data were obtained at the University of Wisconsin – Madison Biophysics Instrumentation Facility, which was established with support from the University of Wisconsin – Madison and grants BIR-9512577 (NSF) and S10 RR13790 (NIH).

## Supplemental Information for <u>Opsins are Scramblases in All Domains of Life</u> Maschmann *et al.,* 2025

**Supplemental Figure 1:**
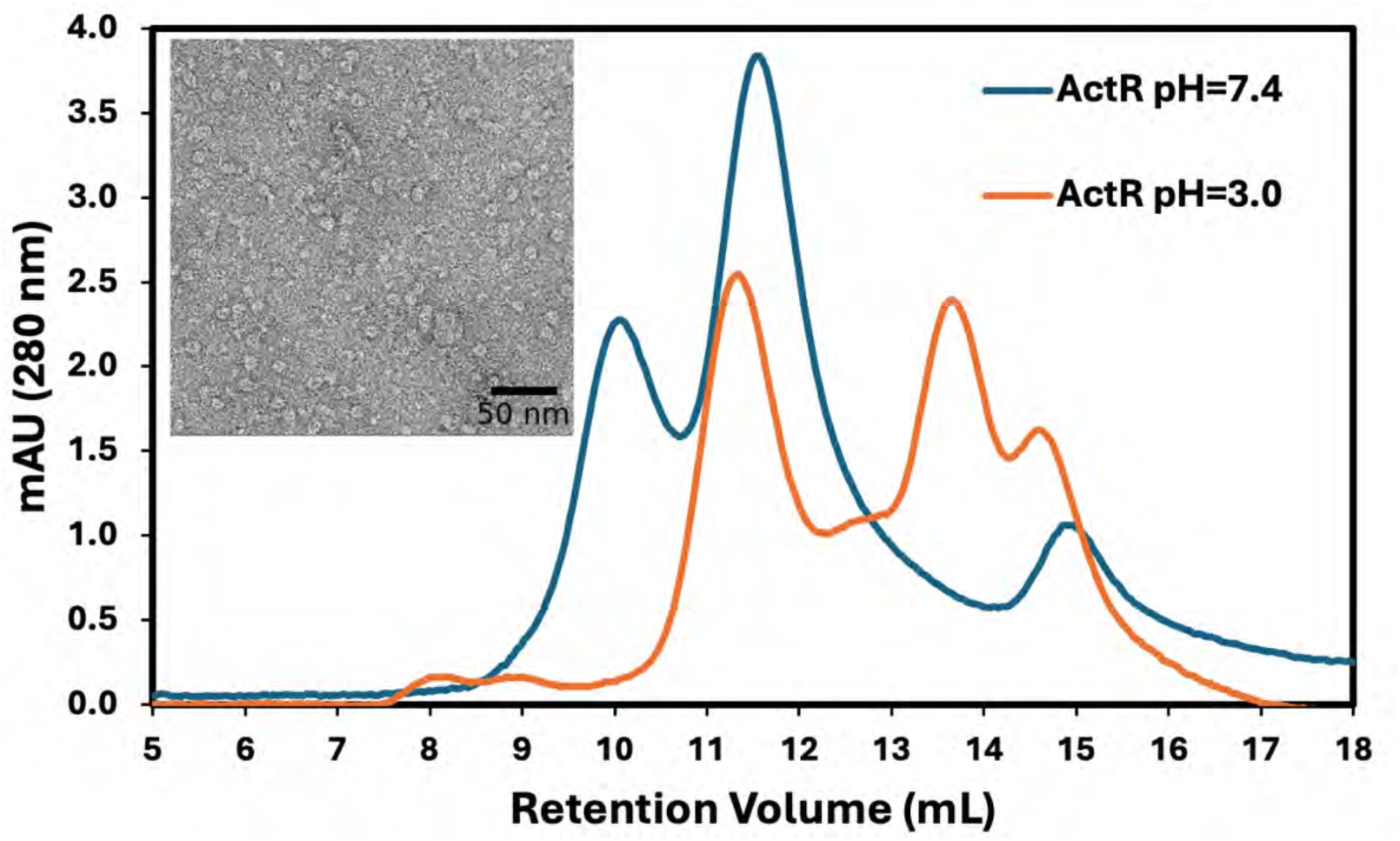
ActR forms Pentamers at Neutral pH. ActR protein purified by nickel a@inity was used for negative stain EM (inset shows grid area, isolated particles shown in Figure 2) and was run on an S200 Increase 10/300 GL SEC column at either pH 7.4 or 3.0. The peak at ∼10 mL in the pH 7.4 trace is consistent with pentameric ActR. The oligomer equilibrium shifts to trimer and monomer at low pH, which is similar to behaviour reported for *Gloeobacter* rhodopsin (1).

### ActR pentamer equilibrium is a function of pH

We investigated the oligomeric state of ActR and possible pH-dependent changes to this assembly using analytical size-exclusion chromatography (SEC). At pH=7.4, ActR predominantly assumes two oligomeric states as resolved by SEC (**Suppl** Figure 2). Based on expected retention volumes of soluble protein standards provided by Cytiva, these SEC peaks are consistent with pentameric and trimeric ActR. Furthermore, ActR changes the distribution of its oligomeric states at pH=3.0 (**Suppl** Figure 2), where the higher molecular weight, presumably pentameric, ActR peak disappears and ActR assumes a primarily trimeric state with a smaller molecular weight peak, possibly monomeric or dimeric ActR. Corollary experiments at pH=5.0 reveal similar SEC traces to those at pH=3.0 (not shown), consistent with the notion that the protonation state of a surface-exposed histidine residue such as H61 is responsible for controlling the stability of states. The mixture of oligomers at both pH=7.4 and pH=3.0 indicate that an equilibrium exists among oligomeric states of ActR. Stable oligomers ordinarily elute from a size-exclusion column in relatively narrow peaks. Contrariwise, ActR at all examined pH values exhibits relatively broad SEC elution peaks, suggesting that ActR oligomers exchange subunits over the course of size-exclusion.

To verify the pentameric state at neutral pH, electron microscopy images were taken of ActR following nickel aXinity chromatography, and indicate a heterogeneous mixture of ActR oligomers (**Suppl** Figure 2). As the protonation state of H61 likely modulates the equilibrium of ActR oligomers with pH, the presence of imidazole in the sample probably eXects the heterogeneity of conformations observed for ActR multimers. Nevertheless, ActR pentamers can be easily observed (Figure 2) and representative negative stain SEM images fit the profile of an AlphaFold2 model of pentameric ActR (Figure 2).

**Supplemental Figure 2:**
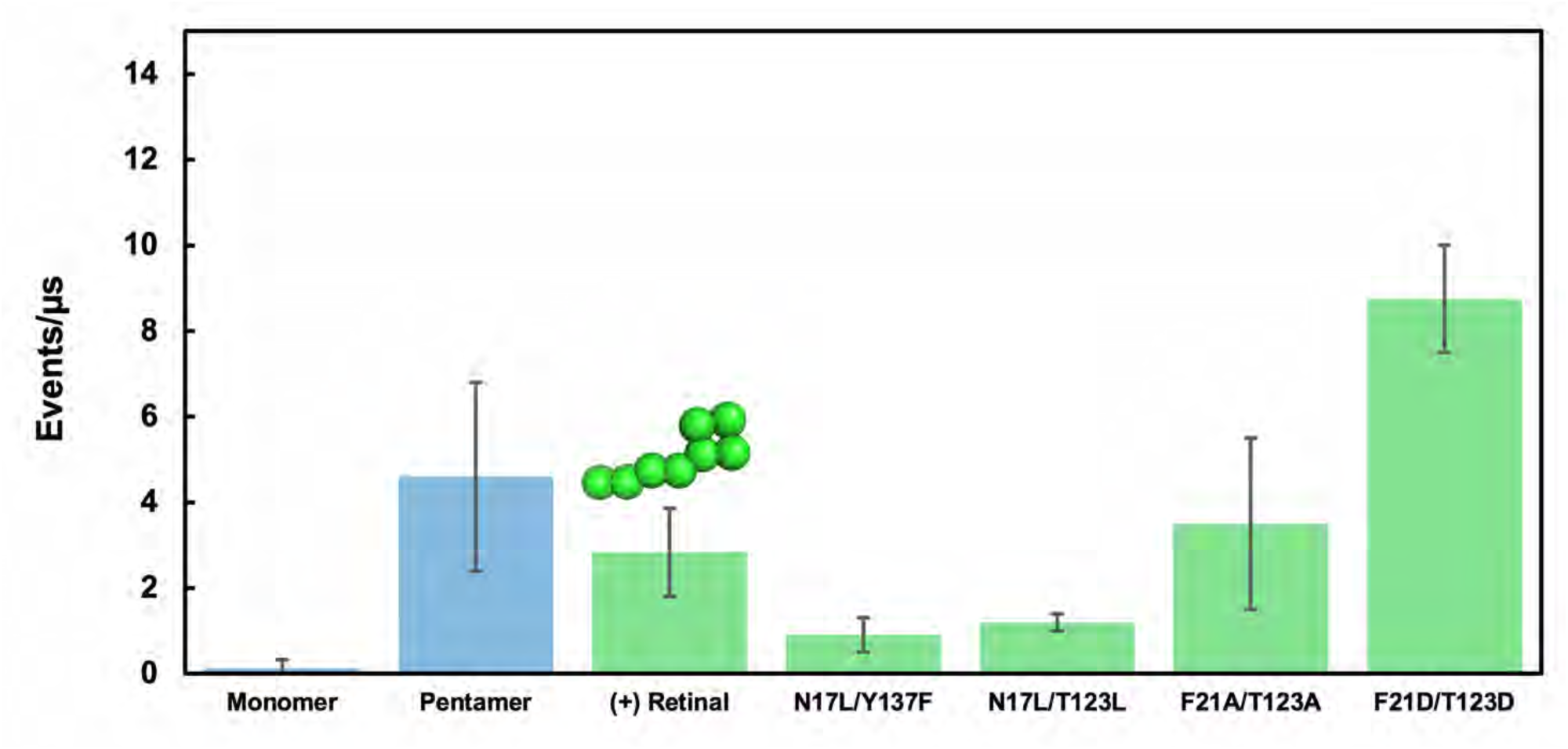
CGMD Scramblase Activity Depends on Polar Residues in Protomer Interface. Monomer and pentamer rates are from Figure 2. The coarse-grained bead model for retinal is represented above the bar for scrambling rates with a retinal-bound opsin.

### CGMD with retinal

A minimized all-atom pentamer model with retinal was used as the basis for positioning the CG ActR model, thus capturing the coordinates in order to subsequently create a GC retinal model. A python script was used to find centroids every 2-3 beads, 4 in the aliphatic tail and 4 for β-ionone ring of retinal. For bead definitions, SC (small) beads correspond to hydrocarbon backbones and TC (tiny) beads are typically used for rings because they allow for rigidity and tighter packing; the “1” is apolar, and the “2” is slightly more polar (maximum polarity is 4). The first bead of the tail is SC2, and the rest are SC1. All the ring beads are TC1. This approach created a simplified, space-filling model of the retinal ligand rather than a rigorous representation of a conjugated C20 ligand. During GROMACS production runs, a “pull-code” was used to keep the retinal ligands in place. This approach uses harmonic restraints between the centers of mass of two atoms to maintain a specified distance between them. Strong forces of 8000 kj/(mol*nm) were applied between bead 1 of the tail and the side chain bead of lysine 233, with a weaker force of 500 kj/(mol*nm) between Gly136 and one side of the ring. Subsequent setup and MD runs and scramblase event counting were carried out as described in the main methods section. Result is the average of 3 x 2 μsec MD simulations.

### CGMD with amino acid substitutions

The variant opsins were created by substituting side chains in PyMol, followed by vacuum minimization and subsequent steps as described in main text except that the membrane contained 800 DOPC molecules rather than 1000. All variants were tested in two independent runs with consistent outputs.

We designed three kinds of amino acid variants. First, in order to directly test the hypothesis that polar residues in the interface were important for interaction with the phospholipid head group we made two models that each had two amino acid substitutions in the scrambling cleft, from polar to non-polar side chains but of similar size. In one case the positions were at the top and bottom of the cleft on opposing subunits (N17L/T123L) and in the other both were at the top of the cleft, again on opposite faces (N17L/Y137F). Both of these reduced scrambling to <1 event/ μsec (**Suppl.** Fig. 2). As a check that not all substitutions would inhibit scrambling, two models were made with modest substitutions to alanine. In a double mutant (F21A/T123A) rates were not changed significantly. In a triple mutant (Y32A/Y93A/Y137A) scrambling was slightly higher; perhaps three substitutions compromise normal oligomer packing. Finally, introduction of two acidic side chains was tested (F21D/T123D), with an increase in scrambling rate. In our simplistic coarse-grained, implicit solvent model we neutralized each of these Asp residues by setting their charge to 0 in the parameter file and using the P2 polar bead type, reasoning that Asp side chains buried within the lipid bilayer have a higher pKa than those exposed to aqueous solvent. 10 Na+ beads were deleted to accommodate and balance the charge of the system. Future research might explore whether the higher apparent pKa of the carboxylic acid side chains within the membrane keeps these polar side chains protonated (uncharged) and, as a consequence, they may not repel the phosphate head groups but pull them through the cleft.

## References

1. Zhang Y-M, Rock CO. 2008. Membrane lipid homeostasis in bacteria. Nat Rev Microbiol 6:222–233.

2. Sanyal S, Menon AK. 2009. Flipping Lipids: Why an’ What’s the Reason for? ACS Chem Biol 4:895–909.

3. McConnell HM, Kornberg RD. 1971. Inside-outside transitions of phospholipids in vesicle membranes. Biochemistry 10:1111–1120.

4. Rothman JE, Kennedy EP. 1977. Rapid transmembrane movement of newly synthesized phospholipids during membrane assembly. Proc Natl Acad Sci U S A 74:1821–1825.

5. Hrafnsdóttir S, Menon AK. 2000. Reconstitution and Partial Characterization of Phospholipid Flippase Activity from Detergent Extracts of the Bacillus subtilis Cell Membrane. J Bacteriol 182:4198–4206.

6. Hrafnsdóttir S, Nichols JW, Menon AK. 1997. Transbilayer Movement of Fluorescent Phospholipids in Bacillus megaterium Membrane Vesicles. Biochemistry 36:4969– 4978.

7. Huijbregts RPH, de Kroon AIPM, de Kruijff B. 1996. Rapid transmembrane movement of C6-NBD-labeled phospholipids across the inner membrane of *Escherichia coli*. Biochim Biophys Acta BBA – Biomembr 1280:41–50.

8. Kubelt J, Menon AK, Müller P, Herrmann A. 2002. Transbilayer Movement of Fluorescent Phospholipid Analogues in the Cytoplasmic Membrane of Escherichia coli. Biochemistry 41:5605–5612.

9. Kol MA, de Kruijff B, de Kroon AIPM. 2002. Phospholipid flip-flop in biogenic membranes: what is needed to connect opposite sides. Semin Cell Dev Biol 13:163– 170.

10. Pomorski TG, Menon AK. 2016. Lipid somersaults: Uncovering the mechanisms of protein-mediated lipid flipping. Prog Lipid Res 64:69–84.

11. Luisi PL, Walde P, Oberholzer T. 1999. Lipid vesicles as possible intermediates in the origin of life. Curr Opin Colloid Interface Sci 4:33–39.

12. Deamer DW. 1997. The first living systems: a bioenergetic perspective. Microbiol Mol Biol Rev 61:239–261.

13. Sebinelli HG, Syska C, Čopič A, Lenoir G. 2024. Established and emerging players in phospholipid scrambling: A structural perspective. Biochimie S0300-9084(24)00218–9.

14. Wang Y, Kinoshita T. 2023. The role of lipid scramblases in regulating lipid distributions at cellular membranes. Biochem Soc Trans 51:1857–1869.

15. Pomorski T, Menon AK. 2006. Lipid flippases and their biological functions. Cell Mol Life Sci CMLS 63:2908–2921.

16. Sakuragi T, Nagata S. 2023. Regulation of phospholipid distribution in the lipid bilayer by flippases and scramblases. Nat Rev Mol Cell Biol 24:576–596.

17. Menon I, Huber T, Sanyal S, Banerjee S, Barré P, Canis S, Warren JD, Hwa J, Sakmar TP, Menon AK. 2011. Opsin Is a Phospholipid Flippase. Curr Biol 21:149–153.

18. Hessel E, Herrmann A, Müller P, Schnetkamp PPM, Hofmann K-P. 2000. The transbilayer distribution of phospholipids in disc membranes is a dynamic equilibrium. Eur J Biochem 267:1473–1483.

19. Wu G, Hubbell WL. 1993. Phospholipid asymmetry and transmembrane diffusion in photoreceptor disc membranes. Biochemistry 32:879–888.

20. Verchère A, Ou W-L, Ploier B, Morizumi T, Goren MA, Bütikofer P, Ernst OP, Khelashvili G, Menon AK. 2017. Light-independent phospholipid scramblase activity of bacteriorhodopsin from Halobacterium salinarum. Sci Rep 7:9522.

21. Beja O, Aravind L, Koonin EV, Suzuki MT, Hadd A, Nguyen LP, Jovanovich SB, Gates CM, Feldman RA, Spudich JL, Spudich EN, DeLong EF. 2000. Bacterial rhodopsin: evidence for a new type of phototrophy in the sea. Science 289:1902–1906.

22. Sharma AK, Zhaxybayeva O, Papke RT, Doolittle WF. 2008. Actinorhodopsins: proteorhodopsin-like gene sequences found predominantly in non-marine environments. Environ Microbiol 10:1039–1056.

23. Dwulit-Smith JR, Hamilton JJ, Stevenson DM, He S, Oyserman BO, Moya-Flores F, Garcia SL, Amador-Noguez D, McMahon KD, Forest KT. 2018. acI Actinobacteria Assemble a Functional Actinorhodopsin with Natively Synthesized Retinal. Appl Environ Microbiol 84:e01678–18.

24. Keffer JL, Hahn MW, Maresca JA. 2016. Characterization of an Unconventional Rhodopsin from the Freshwater Actinobacterium Rhodoluna lacicola. J Bacteriol 197:2704–2712.

25. Nakamura S, Kikukawa T, Tamogami J, Kamiya M, Aizawa T, Hahn MW, Ihara K, Kamo N, Demura M. 2016. Photochemical characterization of actinorhodopsin and its functional existence in the natural host. Biochim Biophys Acta BBA – Bioenerg 1857:1900–1908.

26. Hirschi S, Lemmin T, Ayoub N, Kalbermatter D, Pellegata D, Ucurum Z, Gertsch J, Fotiadis D. 2024. Structural insights into the mechanism and dynamics of proteorhodopsin biogenesis and retinal scavenging. Nat Commun 15:6950.

27. Wu X, Jiang J, Hu J. 2013. Determination and occurrence of retinoids in a eutrophic lake (Taihu Lake, China): cyanobacteria blooms produce teratogenic retinal. Environ Sci Technol 47:807–814.

28. Jaffe AL, Konno M, Kawasaki Y, Kataoka C, Béjà O, Kandori H, Inoue K, Banfield JF. 2022. Saccharibacteria harness light energy using type-1 rhodopsins that may rely on retinal sourced from microbial hosts. ISME J 16:2056–2059.

29. Singh N, Bhalla N. 2020. Moonlighting Proteins. Annu Rev Genet 54:265–285.

30. Hamilton JJ, Garcia SL, Brown BS, Oyserman BO, Moya-Flores F, Bertilsson S, Malmstrom RR, Forest KT, McMahon KD. 2017. Metabolic Network Analysis and Metatranscriptomics Reveal Auxotrophies and Nutrient Sources of the Cosmopolitan Freshwater Microbial Lineage acI. mSystems 2:10.1128/msystems.00091-17.

31. Corcelli A, Lattanzio VMT, Mascolo G, Papadia P, Fanizzi F. 2002. Lipid-protein stoichiometries in a crystalline biological membrane: NMR quantitative analysis of the lipid extract of the purple membrane. J Lipid Res 43:132–140.

32. Fang J, Zhang Y, Zhu T, Li Y. 2023. Scramblase activity of proteorhodopsin confers physiological advantages to Escherichia coli in the absence of light. iScience 26:108551.

33. Aly KA, Beebe ET, Chan CH, Goren MA, Sepúlveda C, Makino S, Fox BG, Forest KT. 2013. Cell-free production of integral membrane aspartic acid proteases reveals zinc-dependent methyltransferase activity of the Pseudomonas aeruginosa prepilin peptidase PilD. MicrobiologyOpen 2:94–104.

34. Mao J, Jin X, Shi M, Heidenreich D, Brown LJ, Brown RCD, Lelli M, He X, Glaubitz C. 2024. Molecular mechanisms and evolutionary robustness of a color switch in proteorhodopsins. Sci Adv 10:eadj0384.

35. Ploier B, Menon AK. 2016. A Fluorescence-based Assay of Phospholipid Scramblase Activity. J Vis Exp JoVE 54635.

36. Goren MA, Morizumi T, Menon I, Joseph JS, Dittman JS, Cherezov V, Stevens RC, Ernst OP, Menon AK. 2014. Constitutive phospholipid scramblase activity of a G Protein-coupled receptor*. Nat Commun 5:5115.

37. Kovalev K, Astashkin R, Gushchin I, Orekhov P, Volkov D, Zinovev E, Marin E, Rulev M, Alekseev A, Royant A, Carpentier P, Vaganova S, Zabelskii D, Baeken C, Sergeev I, Balandin T, Bourenkov G, Carpena X, Boer R, Maliar N, Borshchevskiy V, Büldt G, Bamberg E, Gordeliy V. 2020. Molecular mechanism of light-driven sodium pumping. Nat Commun 11:2137.

38. Morizumi T, Ou W-L, Van Eps N, Inoue K, Kandori H, Brown LS, Ernst OP. 2019. X-ray Crystallographic Structure and Oligomerization of Gloeobacter Rhodopsin. Sci Rep 9:11283.

39. Jahn H, Bartoš L, Dearden GI, Dittman JS, Holthuis JCM, Vácha R, Menon AK. 2023. Phospholipids are imported into mitochondria by VDAC, a dimeric beta barrel scramblase. Nat Commun 14:8115.

40. Iizuka A, Kajimoto K, Fujisawa T, Tsukamoto T, Aizawa T, Kamo N, Jung K-H, Unno M, Demura M, Kikukawa T. 2019. Functional importance of the oligomer formation of the cyanobacterial H+ pump Gloeobacter rhodopsin. Sci Rep 9:10711.

41. Soto-Rodríguez J, Baneyx F. 2019. Role of the signal sequence in proteorhodopsin biogenesis in E. coli. Biotechnol Bioeng 116:912–918.

42. Morizumi T, Ou W-L, Van Eps N, Inoue K, Kandori H, Brown LS, Ernst OP. 2019. X-ray Crystallographic Structure and Oligomerization of Gloeobacter Rhodopsin. Sci Rep 9:11283.

43. Ayoub N, Djabeur N, Harder D, Jeckelmann J-M, Ucurum Z, Hirschi S, Fotiadis D. 2025. Actinorhodopsin: an efficient and robust light-driven proton pump for bionanotechnological applications. Sci Rep 15:4054.

44. Li D, Rocha-Roa C, Schilling MA, Reinisch KM, Vanni S. 2024. Lipid scrambling is a general feature of protein insertases. Proc Natl Acad Sci 121:e2319476121.

45. Morra G, Razavi AM, Pandey K, Weinstein H, Menon AK, Khelashvili G. 2018. Mechanisms of lipid scrambling by the G protein-coupled receptor opsin. Struct Lond Engl 1993 26:356–367.e3.

46. Lee B-C, Menon AK, Accardi A. 2016. The nhTMEM16 Scramblase Is Also a Nonselective Ion Channel. Biophys J 111:1919–1924.

47. Bartoš L, Menon AK, Vácha R. 2024. Insertases scramble lipids: Molecular simulations of MTCH2. Struct Lond Engl 1993 32:505–510.e4.

48. Kalienkova V, Clerico Mosina V, Paulino C. 2021. The Groovy TMEM16 Family: Molecular Mechanisms of Lipid Scrambling and Ion Conduction. 16. J Mol Biol 433:166941.

49. Sephus CD, Fer E, Garcia AK, Adam ZR, Schwieterman EW, Kacar B. 2022. Earliest Photic Zone Niches Probed by Ancestral Microbial Rhodopsins. Mol Biol Evol 39:msac100.

50. DasSarma S, Schwieterman EW. 2021. Early evolution of purple retinal pigments on Earth and implications for exoplanet biosignatures. Int J Astrobiol 20:241–250.

51. Bryant DA, Frigaard N-U. 2006. Prokaryotic photosynthesis and phototrophy illuminated. Trends Microbiol 14:488–496.

52. Pinhassi J, DeLong EF, Béjà O, González JM, Pedrós-Alió C. 2016. Marine Bacterial and Archaeal Ion-Pumping Rhodopsins: Genetic Diversity, Physiology, and Ecology. Microbiol Mol Biol Rev MMBR 80:929–954.

53. Sharma AK, Spudich JL, Doolittle WF. 2006. Microbial rhodopsins: functional versatility and genetic mobility. Trends Microbiol 14:463–469.

54. Hankins MTK, Parrag M, Garaeva AA, Earp JC, Seeger MA, Stansfeld PJ, Bublitz M. 2025. MprF from Pseudomonas aeruginosa is a promiscuous lipid scramblase with broad substrate specificity. Sci Adv 11:eads9135.

55. Arndt M, Alvadia C, Straub MS, Clerico Mosina V, Paulino C, Dutzler R. 2022. Structural basis for the activation of the lipid scramblase TMEM16F. Nat Commun 13:6692.

56. Nakao H, Tsujii T, Saito H, Ikeda K, Nakano M. 2025. Synergistic effects of hydrophilic residues in the transmembrane region on lipid scrambling activity of dimeric helices. Colloids Surf B Biointerfaces 251:114612.

57. Nakao H, Sugimoto Y, Ikeda K, Saito H, Nakano M. 2020. Structural Feature of Lipid Scrambling Model Transmembrane Peptides: Same-Side Positioning of Hydrophilic Residues and Their Deeper Position. J Phys Chem Lett 11:1662–1667.

58. Marsh D. 2013. Handbook of Lipid Bilayers, 2nd ed. CRC Press, Boca Raton.

59. Rohwer RR, Kirkpatrick M, Garcia SL, Kellom M, McMahon KD, Baker BJ. 2025. Two decades of bacterial ecology and evolution in a freshwater lake. Nat Microbiol 10:246– 257.

60. Balashov SP, Imasheva ES, Boichenko VA, Antón J, Wang JM, Lanyi JK. 2005. Xanthorhodopsin: A Proton Pump with a Light-Harvesting Carotenoid Antenna. Science 309:2061–2064.

61. Chuon K, Shim J, Kang K-W, Cho S-G, Hour C, Meas S, Kim J-H, Choi A, Jung K-H. 2022. Carotenoid binding in Gloeobacteria rhodopsin provides insights into divergent evolution of xanthorhodopsin types. Commun Biol 5:1–8.

62. Tzlil G, Marín MDC, Matsuzaki Y, Nag P, Itakura S, Mizuno Y, Murakoshi S, Tanaka T, Larom S, Konno M, Abe-Yoshizumi R, Molina-Márquez A, Bárcenas-Pérez D, Cheel J, Koblížek M, León R, Katayama K, Kandori H, Schapiro I, Shihoya W, Nureki O, Inoue K, Rozenberg A, Chazan A, Béjà O. 2025. Structural insights into light harvesting by antenna-containing rhodopsins in marine Asgard archaea. Nat Microbiol 10:1484– 1500.

63. Garcia SL, McMahon KD, Martinez-Garcia M, Srivastava A, Sczyrba A, Stepanauskas R, Grossart HP, Woyke T, Warnecke F. 2013. Metabolic potential of a single cell belonging to one of the most abundant lineages in freshwater bacterioplankton. ISME J 7:137– 147.

64. Gibson DG, Young L, Chuang R-Y, Venter JC, Hutchison CA, Smith HO. 2009. Enzymatic assembly of DNA molecules up to several hundred kilobases. Nat Methods 6:343– 345.

65. Heydenreich FM, Miljuš T, Jaussi R, Benoit R, Milić D, Veprintsev DB. 2017. High-throughput mutagenesis using a two-fragment PCR approach. Sci Rep 7:6787.

66. Beebe ET, Makino S-I, Nozawa A, Matsubara Y, Frederick RO, Primm JG, Goren MA, Fox BG. 2011. Robotic large-scale application of wheat cell-free translation to structural studies including membrane proteins. New Biotechnol 28:239–249.

67. Dearden GI, Ravishankar V, Sakata K, Menon AK, Bergdoll L. 2024. Protocol for the production and reconstitution of VDAC1 for functional assays. STAR Protoc 5:103240.

68. Ong SGM, Chitneni M, Lee KS, Ming LC, Yuen KH. 2016. Evaluation of Extrusion Technique for Nanosizing Liposomes. 4. Pharmaceutics 8:36.

69. Brunner JD, Schenck S, Dutzler R. 2016. Structural basis for phospholipid scrambling in the TMEM16 family. Curr Opin Struct Biol 39:61–70.

70. Malvezzi M, Chalat M, Janjusevic R, Picollo A, Terashima H, Menon AK, Accardi A. 2013. Ca2+-dependent phospholipid scrambling by a reconstituted TMEM16 ion channel. Nat Commun 4:2367.

71. McIntyre JC, Sleight RG. 1991. Fluorescence assay for phospholipid membrane asymmetry. Biochemistry 30:11819–11827.

72. Chang Q, Gummadi SN, Menon AK. 2004. Chemical Modification Identifies Two Populations of Glycerophospholipid Flippase in Rat Liver ER. Biochemistry 43:10710– 10718.

73. Menon I, Sych T, Son Y, Morizumi T, Lee J, Ernst OP, Khelashvili G, Sezgin E, Levitz J, Menon AK. 2024. A cholesterol switch controls phospholipid scrambling by G protein-coupled receptors. J Biol Chem 300:105649.

74. Team CD, Boitreaud J, Dent J, McPartlon M, Meier J, Reis V, Rogozhonikov A, Wu K. 2024. Chai-1: Decoding the molecular interactions of life. bioRxiv 10.1101/2024.10.10.615955.

75. Razeto A, Gribbon P, Loew C. 2019. The dynamic nature of the VDAC1 channels in bilayers: human VDAC1 at 2.7 Angstrom resolution: 6g6u.

76. Abramson J, Adler J, Dunger J, Evans R, Green T, Pritzel A, Ronneberger O, Willmore L, Ballard AJ, Bambrick J, Bodenstein SW, Evans DA, Hung C-C, O’Neill M, Reiman D, Tunyasuvunakool K, Wu Z, Žemgulytė A, Arvaniti E, Beattie C, Bertolli O, Bridgland A, Cherepanov A, Congreve M, Cowen-Rivers AI, Cowie A, Figurnov M, Fuchs FB, Gladman H, Jain R, Khan YA, Low CMR, Perlin K, Potapenko A, Savy P, Singh S, Stecula A, Thillaisundaram A, Tong C, Yakneen S, Zhong ED, Zielinski M, Žídek A, Bapst V, Kohli P, Jaderberg M, Hassabis D, Jumper JM. 2024. Accurate structure prediction of biomolecular interactions with AlphaFold 3. Nature 630:493–500.

77. Bai L, You Q, Feng X, Kovach A, Li H. 2020. Structure of the ER membrane complex, a transmembrane-domain insertase. Nature 584:475–478.

78. Bekker H, Berendsen HJC, Djikstra EJ, Achterop S, van Drunen R, van der Spoel D, Sijbers A, Keegstra H. 1993. Gromacs: A parallel computer for molecular dynamics simulations, p. 252–256. *In* de Groot, RA, Nadrchal, J (eds.), Physics Computing’ 92. World Scientific, Singapore.

79. Thain D, Tannenbaum T, Livny M. 2005. Distributed computing in practice: the Condor experience. Concurr – Pract Exp 17:323–356.

80. Humphrey W, Dalke A, Schulten K. 1996. VMD: Visual molecular dynamics. J Mol Graph 14:33–38.

81. Stone J. 1998. An Efficient Library for Parallel Ray Tracing and Animation. Computer Science Department, University of Missouri-Rolla.

82. Kimura Y, Vassylyev DG, Miyazawa A, Kidera A, Matsushima M, Mitsuoka K, Murata K, Hirai T, Fujiyoshi Y. 1997. Surface of bacteriorhodopsin revealed by high-resolution electron crystallography. Nature 389:206–211.

83. Lomize MA, Pogozheva ID, Joo H, Mosberg HI, Lomize AL. 2012. OPM database and PPM web server: resources for positioning of proteins in membranes. Nucleic Acids Res 40:D370–376.

84. Jo S, Kim T, Iyer VG, Im W. 2008. CHARMM-GUI: A web-based graphical user interface for CHARMM. J Comput Chem 29:1859–1865.

85. Brooks BR, Brooks III CL, Mackerell Jr. AD, Nilsson L, Petrella RJ, Roux B, Won Y, Archontis G, Bartels C, Boresch S, Caflisch A, Caves L, Cui Q, Dinner AR, Feig M, Fischer S, Gao J, Hodoscek M, Im W, Kuczera K, Lazaridis T, Ma J, Ovchinnikov V, Paci E, Pastor RW, Post CB, Pu JZ, Schaefer M, Tidor B, Venable RM, Woodcock HL, Wu X, Yang W, York DM, Karplus M. 2009. CHARMM: The biomolecular simulation program. J Comput Chem 30:1545–1614.

86. Lee J, Cheng X, Swails JM, Yeom MS, Eastman PK, Lemkul JA, Wei S, Buckner J, Jeong JC, Qi Y, Jo S, Pande VS, Case DA, Brooks CLI, MacKerell AD Jr, Klauda JB, Im W. 2016. CHARMM-GUI Input Generator for NAMD, GROMACS, AMBER, OpenMM, and CHARMM/OpenMM Simulations Using the CHARMM36 Additive Force Field. J Chem Theory Comput 12:405–413.

87. Qi Y, Ingólfsson HI, Cheng X, Lee J, Marrink SJ, Im W. 2015. CHARMM-GUI Martini Maker for Coarse-Grained Simulations with the Martini Force Field. J Chem Theory Comput 11:4486–4494.

88. Hsu P-C, Bruininks BMH, Jefferies D, Cesar Telles de Souza P, Lee J, Patel DS, Marrink SJ, Qi Y, Khalid S, Im W. 2017. CHARMM-GUI Martini Maker for modeling and simulation of complex bacterial membranes with lipopolysaccharides. J Comput Chem 38:2354–2363.

89. Center for High Throughput Computing. (2006). doi:10.21231/GNT1-HW21

## Reference

1. Morizumi T, Ou W-L, Van Eps N, Inoue K, Kandori H, Brown LS, Ernst OP. 2019. X-ray Crystallographic Structure and Oligomerization of Gloeobacter Rhodopsin. Sci Rep 9:11283.

